# Fmrp regulates neuronal balance in embryonic motor circuit formation

**DOI:** 10.1101/2022.06.06.495019

**Authors:** Chase M. Barker, Kaleb D. Miles, Caleb A. Doll

**Author notes:** Author Contributions C.M.B.: Software, Formal Analysis, Investigation. K.D.M.: Investigation. C.A.D: Conceptualization, Formal Analysis, Investigation, Writing, Visualization.

## Abstract

Motor behavior requires the balanced production and integration of a variety of neural cell types. Motor neurons are positioned in discrete locations in the ventral spinal cord, targeting specific muscles to drive locomotive contractions. Specialized spinal interneurons modulate and synchronize motor neuron activity to achieve coordinated motor output. Changes in the ratios of spinal interneurons could drastically alter motor output by tipping the balance of inhibition and excitation onto target motor neurons. Importantly, individuals with Fragile X syndrome (FXS) and associated autism spectrum disorders often have significant motor challenges, including repetitive behaviors and epilepsy. FXS stems from the transcriptional silencing of the gene Fragile X Messenger Ribonucleoprotein 1 (FMR1), which encodes an RNA binding protein that is implicated in a multitude of crucial neurodevelopmental processes, including cell specification. We find that zebrafish *fmr1* mutants generate surplus ventral lateral descending (VeLD) interneurons, an early-born cell derived from the pMN domain. These GABAergic interneurons are also associated with changes in synaptogenesis, as *fmr1* mutants show increased early expression of the scaffold Gephyrin, but these postsynaptic sites fail to mature. Our work shows that Fmrp regulates the proportionate production of neurons that comprise early embryonic motor circuits. As VeLD interneurons are hypothesized to act as central pattern generators driving the earliest spontaneous movements, this imbalance could profoundly influence the formation and long-term function of motor circuits driving locomotion.

## Materials and Methods

### Zebrafish lines and husbandry

The Institutional Animal Care and Use Committee at the University of Colorado School of Medicine approved all animal work, which follows the US National Research Council’s Guide for the Care and Use of Laboratory Animals, the US Public Health Service’s Policy on Humane Care and Use of Laboratory Animals, and Guide for the Care and Use of Laboratory Animals. Larvae were raised at 28.5°C in embryo medium and staged as hours (hpf) according to morphological criteria (Kimmel et al., 1995). Zebrafish lines used in this study included *fmr1*^*hu2787*^ (den Broeder et al., 2009), *Tg(olig2:EGFP)*^*vu12*^ (Shin et al., 2003), *Tg(slc17a6b:EGFP)*^*zf139*^ (Miyasaka et al., 2009), *Tg(mnx1:EGFP)*^*ml2*^ (Flanagan-Steet et al., 2005). Genotyping for *fmr1*^*hu2787*^ was performed as previously described (Ng et al., 2013). As zebrafish sex determination does not occur until juvenile stages (Uchida et al., 2002), we were unable to determine the sex of the embryos in our experiments.

### Imaging and Analysis

We acquired images on a Zeiss LSM 880 or a Zeiss CellObserver SD 25 spinning disk confocal system (Carl Zeiss). Images were captured with Zen software (Carl Zeiss), then processed and analyzed using Fiji/ImageJ.

### Fluorescent in situ RNA Hybridization (FISH)

The probes for zebrafish *gata3* and *lhx3* were designed and synthesized by the manufacturer for use with the B2 and B3 amplifiers and B2-546 nm and B3-647 fluorophores, respectively (Molecular Instruments). FISH procedure was guided by the in situ hybridization chain reaction protocol for whole-mount zebrafish (Molecular Instruments, v3.0; Choi et al., 2018). Embryos/larvae at indicated timepoints were fixed in 1 mL of 4% paraformaldehyde (PFA) / 1xPBS for 24 hours at 4°C. Samples underwent 3×5-minute washes with 1 mL of 1×phosphate-buffered saline (PBS) to stop the fixation, followed by 1 mL 4×10-minute and 1×50-minute 100% MeOH washes. Samples were stored overnight at -20°C, then transferred to a 1.5 mL Eppendorf tube and then proceeded through a series of 1 mL MeOH / 0.1% Tween 20/1xPBS (PBSTw) washes for 5 minutes each at room temperature as follows. 1×75% MeOH / 25% PBSTw, 1×50% MeOH / 50% PBSTw, 1×25% MeOH / 75% PBSTw, and 5×100% PBSTw. Samples were treated with Proteinase K (24 hpf at 1:1000 for 5 minutes; 48 hpf at 1:200 for 8 minutes), immediately washed twice with PBSTw (1 mL each) without incubation, postfixed with 1 mL of 4% PFA for 20 minutes at room temperature on a rocker, then washed for 5×5-minutes with 1 mL of PBST. For the detection stage, 500 *µ*L of pre-warmed probe hybridization buffer was added for 30 minutes at 37°C. Then the pre-hybridization solution was removed and 500uL of probe solution (2 uL of probe / 500uL of pre-warmed probe hybridization buffer) was added and incubated overnight (12–16 hrs) at 37°C. The next day, probe wash buffer was warmed to 37°C before starting washes. Samples underwent 4×15-minute washes with 500 μL of probe wash buffer at 37°C followed by 2×5-minute washes with 5×SSCT at room temperature. Before the next wash, the amplification buffer was equilibrated to room temperature, then samples were pre-amplified with 500 μL of amplification buffer for 30 minutes at room temperature. During the pre-amplification step, 30 ρmol of hairpin h1 and 30 ρmol of hairpin h2 were separately prepared by snap cooling 10 μL of 3 μM stock at 95°C for 90 seconds, and then allowed to cool to room temperature in a dark drawer for 30 minutes. The hairpin solution was then prepared by adding snap-cooled h1 hairpins and snap-cooled h2 hairpins to 500 *µ*L of amplification buffer at room temperature. After the completion of the pre-amplification step, the pre-amplification solution was removed, 125 *µ*L of hairpin solution was added, per 1.5 mL tube containing ∼12 embryos/larvae, which were incubated overnight (12–16 h) in the dark at room temperature. The following day the excess hairpins were removed by a series of washes with 500 µL of 5×SSCT for 2×5-minutes, 2×30-minutes, and 1×5-minutes at room temperature. Samples were post-fixed in 1 mL of 4% paraformaldehyde/1xPBS for 20 minutes on a rocker. Then two washes were performed with 1 mL of 1xPBS DEPC with no incubation time.

### Immunohistochemistry

Embryos were fixed in 4% paraformaldehyde/1xPBS and rocked overnight at 4°C. Samples were rinsed in 1xPBS, then embedded in 1.5% agar/30% sucrose blocks and immersed in 30% sucrose overnight. Blocks were frozen on dry ice, then 20 μm transverse sections were taken with a cryostat microtome and collected on polarized slides. Slides were mounted in Sequenza racks (Thermo Scientific), washed twice for 5 minutes in 0.1%Triton/1xPBS (PBSTx) to rehydrate samples, blocked 1 hour in 2%goat serum/2%bovine serum albumin/PBSTx, 200uL per slide. Samples were then incubated in primary antibodies: rabbit α-GABA (1:500; Sigma, A2052; Pedroni and Ampatzis, 2019); mouse α-Gephyrin (1:500; Synaptic Systems; Marisca et al., 2020). Sections were washed 6×15-minutes in PBSTx, then incubated 2 hours at room temperature in 200uL of secondary antibody (1:500; in block): Alexa Fluor goat α-rabbit 488 (A-11008; Invitrogen), Alexa Fluor goat α-mouse 568 (A-11004; Invitrogen). Sections were washed for 1 hour (4×15-minute washes) in PBSTx, incubated with 200 uL of DAPI (1:1000 in PBSTx) for 5 minutes, washed for 30 minutes (2×15-minute washes) in PBSTx, then mounted in Vectashield (Vector Laboratories, H-1000-10). For FISH/IHC combination experiments, wholemount embryos immediately underwent the IHC protocol after hairpin washes and post-fixation.

### Quantification and statistical analyses

For all cell counts, the investigator was blind to genotype. For IHC, DAPI and GABA/Islet channels were used to confirm cell number in each section. For FISH, DAPI and *gata3* or *lhx3* were used to confirm cell number in acquired z-stacks. For quantification of sectioned embryos, values from each group of sections from an individual embryo were averaged (2-4 sections for each sample). All statistics were performed in Graphpad Prism (version 9). Normality was assessed with a D’Agostino and Pearson omnibus test. For two groups, unpaired comparisons were made using either unpaired two-tailed t tests (for normal distributions) or Mann–Whitney tests (abnormal distributions).

### Gephyrin puncta quantification

Gephyrin puncta were quantified from z-projections collected at identical exposures with an ImageJ custom script written by Chase M. Barker: https://github.com/barkerch/Fmrp-regulates-neuronal-balance-in-embryonic-motor-circuit-formation, which is based on a previous method (Scott et al., 2020). First, the center 12 z-intervals of 0.5 μm depth were z-projected using the Standard Deviation projection parameter in ImageJ. Next, the image background was subtracted with a two-rolling ball. Next, the image was thresholded by taking two standard deviations above the mean fluorescence intensity. Two regions of interest (ROIs) were drawn around the entire spinal cord and the ventral spinal cord, and puncta were analyzed using the Analyze Particles feature with a size of 0-Infinity and circularity of 0.00-1.00. ROI areas were then measured using the Measure feature, collecting the quantity and average size (area, μm^2^) of puncta in each ROI.

## Introduction

The integration of distinct cell types into neural circuits occurs during precise developmental windows. In zebrafish locomotive circuits, early-born primary motor neurons (MNs) and interneurons (INs) connect to form rudimentary networks that drive early spontaneous behavior (Saint-Amant and Drapeau, 2001). INs are crucial for this process, as isolated MN clusters must develop synchrony with adjacent MN groups to achieve coordinated locomotion by driving muscle contractions throughout the spine (Cazalets et al., 1992; Tresch and Kiehn, 2000). In more mature circuits, spinal INs provide excitatory and inhibitory modulation of motor neuron output to drive refined locomotion (McLean et al., 2007, 2008; Kimura et al., 2013; Callahan et al., 2019). Less is known about IN function in embryogenesis, though some INs appear to function as central pattern generators, synchronizing MN output in adjacent hemisegments to drive early spontaneous behavior (Saint-Amant and Drapeau, 2000, 2001; Warp et al., 2012). Importantly, changes in the relative amount of synchronizing IN output could profoundly influence nascent neural circuit formation and long-term motor function.

Fragile X syndrome (FXS) is the most common heritable cause of intellectual disability, and individuals with FXS commonly have symptoms consistent with hyperexcitable motor behavior, including repetitive movements and epilepsy (Berry-Kravis, 2002; Oakes et al., 2016). Many vertebrate models of FXS show hyperexcitable motor behavior, including zebrafish *fmr1* mutants at both larval and adult stages (Kim et al., 2014; Shamay-Ramot et al., 2015). Although FXS appears to be rooted in neurodevelopmental mechanisms, the bulk of FXS research has focused on synaptic dysfunction in established neural circuits. It has been difficult to pinpoint the pathogenesis of FXS due to the widespread influence of Fragile X Messenger Ribonucleoprotein (FMRP; Fmrp in zebrafish) on synapse formation and function (Huber et al., 2002; Todd et al., 2003; Dictenberg et al., 2008; Doll et al., 2017). A leading hypothesis is that FXS symptoms are rooted in altered excitatory and inhibitory balance, including diminished inhibitory gamma-amino butyric acid (GABA) signaling in mature FXS circuits (Fatemi et al., 2009; Hashemi et al., 2017; Goel et al., 2018). However, there is also evidence that GABAergic signaling in embryogenesis is altered in the absence of FMRP, such that GABA remains depolarizing at later developmental stages (He et al., 2014; Zhang et al., 2022). It is not yet known when motor defects first develop in FXS; we hypothesize that Fmrp is required in early embryogenesis to proportionally specify interneuron subtypes that provide balanced excitation and inhibition in developing locomotive circuits..

There could be many mechanisms leading to disrupted E/I balance in FXS; however, the proportionate production of neuronal cell types driving excitation and inhibition during development likely provides a foundational influence. The specification of distinct cell types from common progenitor domains requires the regulation of unique gene expression profiles that are tailored to a specific lineage. FMRP regulates a diverse catalog of genes and is classically associated with synaptogenesis and neural plasticity (Dictenberg et al., 2008; Darnell et al., 2011; Deng et al., 2013; Doll and Broadie, 2015; Doll et al., 2017; Maurin et al., 2018). Fmrp also regulates cell fate decisions in embryogenesis: for example, Fmrp regulates the balance of cells generated from the pMN domain in the ventral spinal cord, including MNs and oligodendrocyte lineage cells (Doll et al., 2021). pMN progenitors also produce specialized spinal interneurons that modulate MN output to drive muscle contractions (Park et al., 2004; Warp et al., 2012; Svara et al., 2018). We find that zebrafish *fmr1* mutants generate excess GABAergic INs in the ventral spinal cord and show altered synaptogenesis in embryonic stages, at a developmental stage when motor behavior initiates. Fate mapping reveals excess INs are early-born ventral lateral descending (VeLD) cells born in the pMN domain alongside primary motor neurons. Given proposed roles for VeLD neurons as central pattern generators driving the earliest spontaneous contractions (Warp et al., 2012), our work presents a new hypothesis to help explain the origins of hyperexcitability in FXS.

## Results

### Fmrp restricts the production of ventrolateral GABAergic interneurons and the postsynaptic scaffold Gephryin

We previously showed that Fmrp regulates the proportional production of two essential cell types in the ventral spinal cord: oligodendrocyte lineage cells and MNs (Doll et al., 2021). Given changes in GABAergic neurotransmission in FXS (El Idrissi et al., 2005; Kratovac and Corbin, 2013), we next examined gamma-aminobutyric acid (GABA) INs, which provide crucial input to MNs to modulate muscle contractions and achieve coordinated locomotion (Saint-Amant and Drapeau, 2001; Higashijima et al., 2004b; Callahan et al., 2019). We used immunohistochemistry (IHC) on transverse trunk spinal cord sections to detect embryonic GABAergic INs, finding two main classes of cells in the ventral spinal cord: 1) brightly labeled Kolmer-Agduhr (KA) INs that innervate the central canal and are associated with chemosensory function (asterisks, Fig. 1A,B,E,F; Andrzejczuk et al., 2018; Seredick et al., 2014); 2) dimly labeled GABA^+^ cells in the ventrolateral cord (arrows, Fig. 1A,B,E,F). There was a 33% increase in ventrolateral GABA^+^ INs in *fmr1* embryos (Fig. 1C,G) but no change in KA IN quantify at both 24 and 48 hpf (Fig. 1D,H).

**Figure 1.**
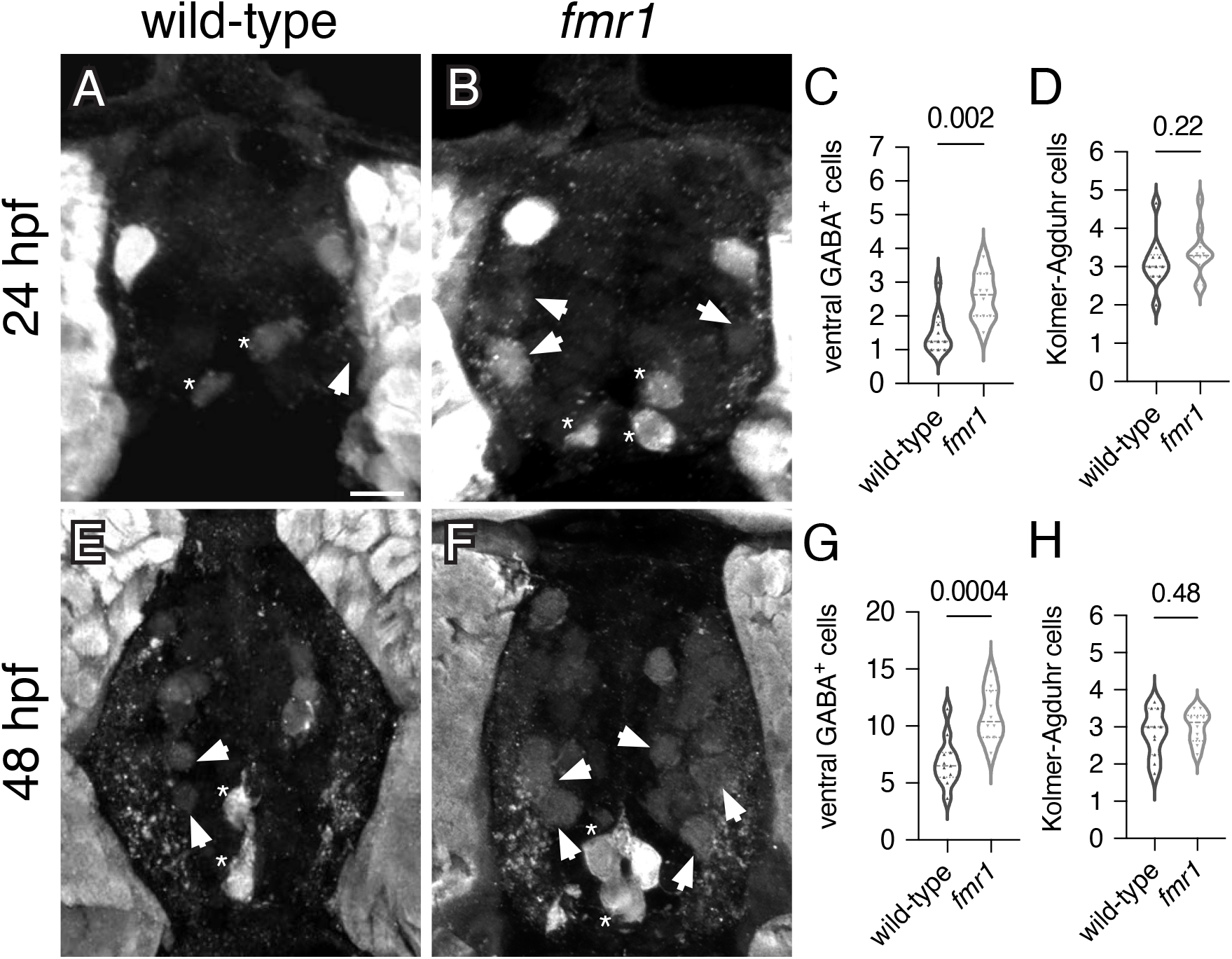
Fmrp restricts the production of GABAergic cells in the ventral spinal cord. Immunohistochemistry to detect GABA at 24 hours post-fertilization (hpf; A,B) and 48 hpf (E,F) on transverse sections of trunk spinal cord reveals two populations of ventral GABA+ interneurons (INs): robustly GABA-expressing Kolmer-Agduhr (KA) INs that line the central canal (asterisks) and a group of more weakly-expressing GABA^+^ INs positioned in the ventrolateral spinal cord (arrowheads). Quantification of ventral GABA^+^ cells at 24 hpf (C) and 24 hpf (C) and 48 hpf (G), and KA neurons at 24 hpf (D) and 48 hpf (H). Scale bar = 10 μm.

As Fmrp also plays crucial roles in the differentiation of neurons and glia (Luo et al., 2010; Edens et al., 2019; Doll et al., 2021; Raj et al., 2021), it is possible that the excess GABAergic cells in *fmr1* mutants are not terminally differentiated and do not form synaptic connections. To address this possibility, we examined expression of the postsynaptic scaffold Gephyrin, which is crucial for the clustering and stabilization of GABAergic and glycinergic receptors (Essrich et al., 1998; Feng et al., 1998). We detected Gephyrin on transverse sections of 24 and 48 hpf trunk spinal cord (Marisca et al., 2020), finding punctate expression in cells lining the central canal and at the ventrolateral edges of the cord, the respective locations of neural progenitors and motor neurons (see insets, Fig. 2A,B,E,F). We quantified Gephyrin puncta and normalized to the relative area of the spinal cord, finding an 18% increase in puncta per unit area in *fmr1* embryos at 24 hpf (Fig. 2C). We also measured Gephyrin puncta size, finding no change in *fmr1* compared with control at this stage (Fig. 2D). At 48 hpf, although the quantity of Gephyrin puncta was no longer elevated in *fmr1* (Fig. 2G), the relative size of Gephyrin puncta in wild-type was 24% larger than in *fmr1* embryos (Fig. 2H). Previous in vitro work demonstrated that Gephyrin puncta size increases over time, such that large puncta are associated with synaptic sites (Craig et al., 1996). This suggests that GABAergic and/or glycinergic postsynaptic maturation in *fmr1* mutant embryos is hindered during this stage of embryonic spinal development. Taken together, our data shows *fmr1* embryos have surplus GABAergic INs in the ventrolateral spinal cord and altered Gephyrin scaffold expression.

**Figure 2.**
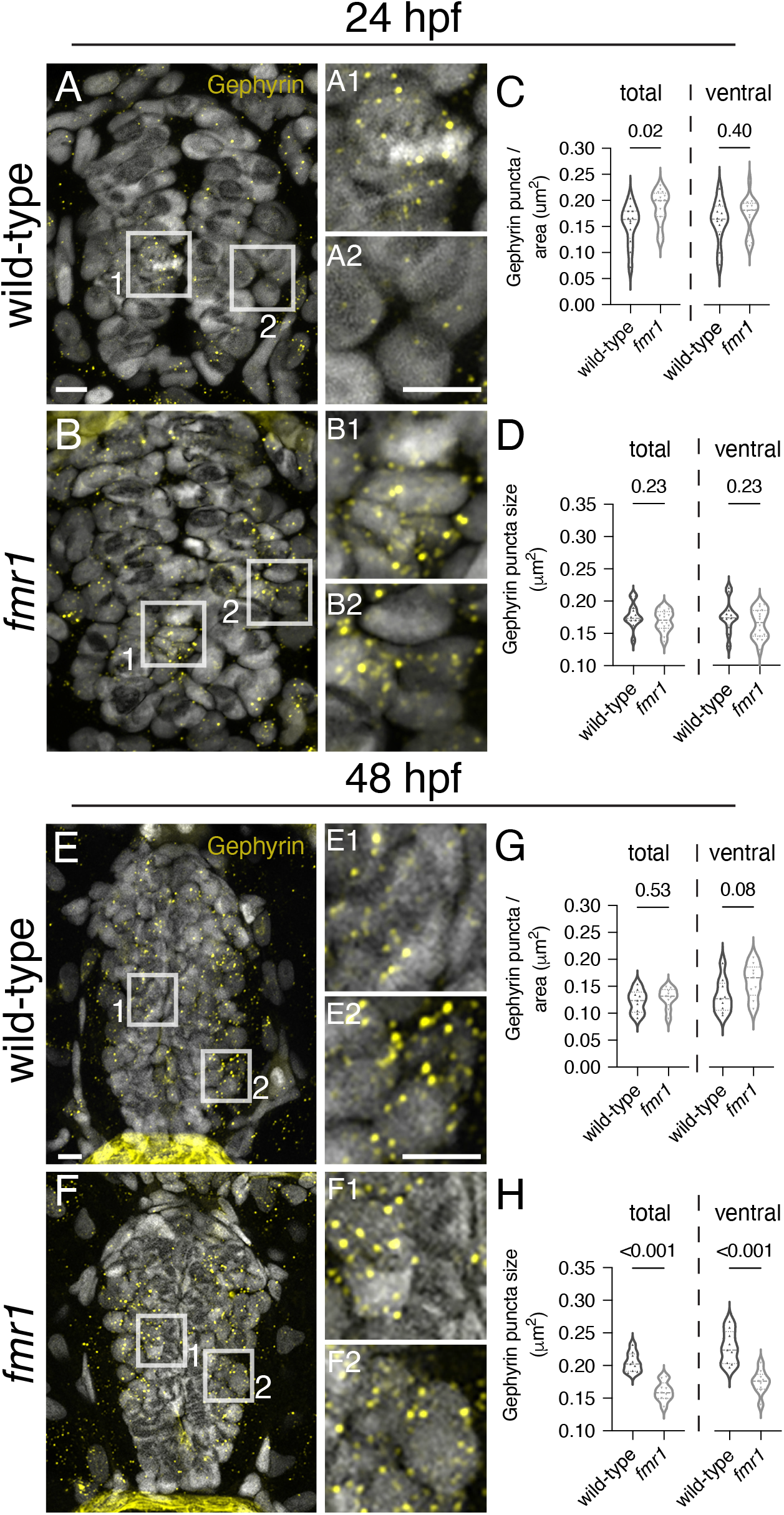
FMRP regulates the production of the postsynaptic scaffold Gephyrin. Representative images from immunohistochemistry experiments to detect Gephyrin on transverse trunk spinal cord sections at 24 hpf (A-B) and 48 hpf (E-F). Gephyrin expression is concentrated in cells adjacent to the central canal (insets A1,B1,E1,F1) and in cells at the ventrolateral edge of the gray matter (insets A2,B2,E2,F2). Quantification of Gephyrin puncta normalized to relative spinal cord area at 24 hpf (C) and 48 hpf (G), in both the total and ventral spinal cord. Quantification of the average size of Gephyrin puncta in wild-type and fmr1 embryos at 24 hpf (D) and 48 hpf (H). Scale bars = 10 μm.

### Production of glutamatergic INs is unchanged in fmr1 mutants

Given the reduction in neural patterning we previously showed in *fmr1* embryos (Doll et al., 2021), it is possible that Fmrp regulates the specification of additional spinal interneurons. Glutamate is the principal excitatory neurotransmitter in the mature spinal cord (Kudo and Yamada, 1987), and glutamatergic INs are specified from several spinal progenitor domains (Higashijima et al., 2004b; Park et al., 2004). To broadly test whether Fmrp also regulates glutamatergic cell production, we used the transgenic reporter *Tg(slc17a6b:EGFP)*, as *slc17a6b* encodes the vesicular glutamate transporter (Miyasaka et al., 2009). *slc17a6b* is expressed two distinct groups of cells in the spinal cord at 24 hpf, including large clusters of cells with small somata in the ventral cord and large Rohon-Beard sensory neurons in the dorsal region (Fig. 3A,B; Higashijima et al., 2004a). There was no change in the average number of EGFP^+^ ventral cells or Rohan-Beard neurons in *fmr1* mutants compared to wild-type control (Fig. 3C-E), which suggests that Fmrp does not regulate the overall production of glutamatergic INs.

**Figure 3.**
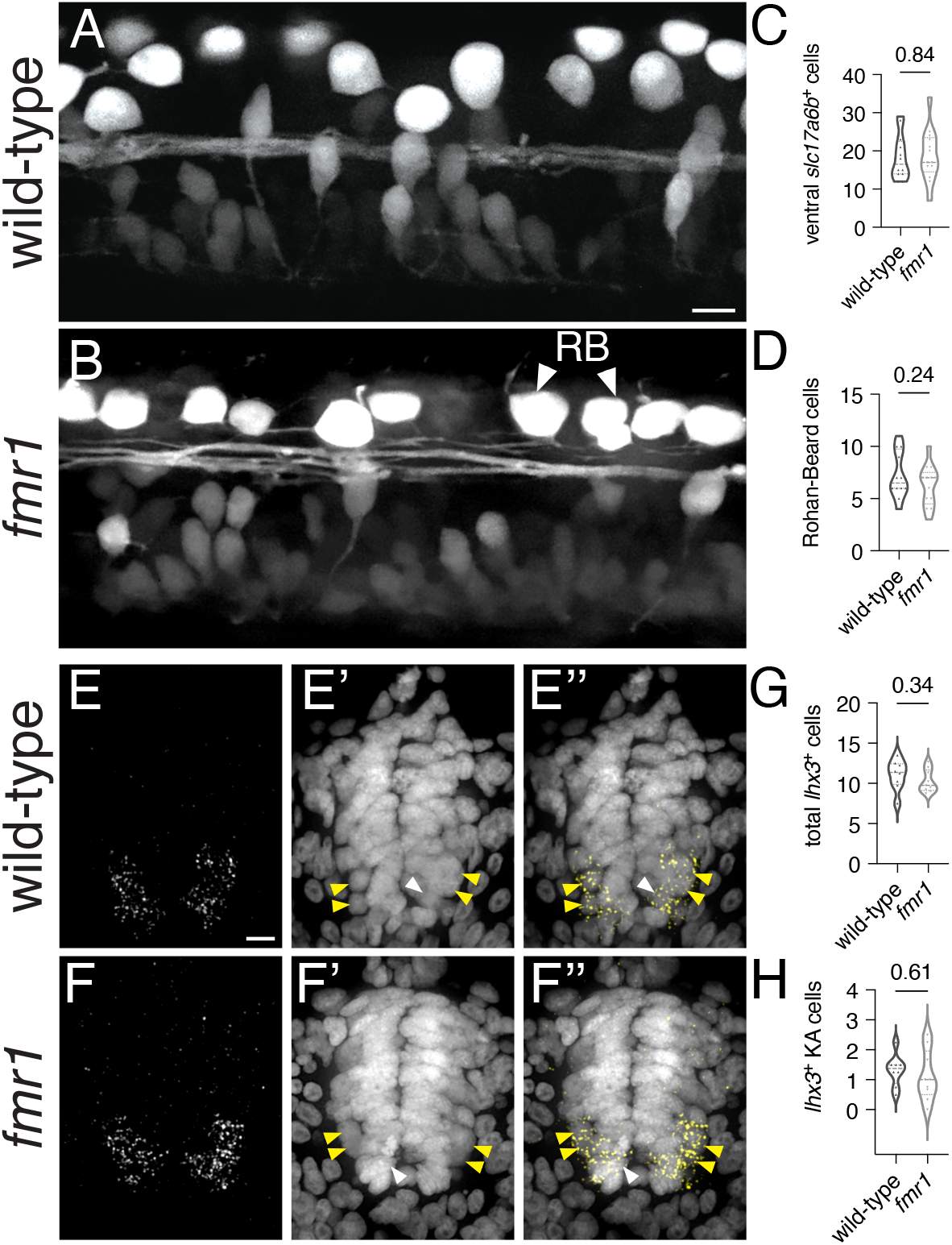
Fmrp does not regulate glutamatergic cell production. Representative lateral images of the spinal cord of live wild-type (A) and *fmr1* mutant (B) embryos expressing *slc17a6b*:LoxP-DsRed-LoxP-EGFP, a glutamatergic neuron reporter, at 24 hpf. Quantification of *slc17a6b*^+^ cells in the ventral spinal cord (C), and *slc17a6b*+ Rohan-Beard (RB) cells in the dorsal cord (D). RBs have large cell bodies and brightly express the reporter (arrowheads, B). Representative images of FISH experiments to detect *Ihx3* expression in transverse trunk spinal cord sections of wildtype (E) and fmr1 (F) embryos at 24 hpf. Presumptive *Ihx3*^+^ V2a I Ns in the lateral region are indicated with yellow arrowheads and *Ihx3*^+^ KA neurons adjacent the central canal are labeled by white arrowheads (E’,F’,E”,F”). Quantification of total *Ihx3*^+^ cells (excluding KA; G) and *Ihx3*^+^ KA cells per section (H). Scale bars = 10 μm.

Embryonic spinal neuron subtypes are defined by unique genetic profiles that are initiated in specific progenitor domains (Park et al., 2004; Seredick et al., 2014; Andrzejczuk et al., 2018). We also examined a specific population of glutamatergic INs in the ventral spinal cord that are specified in the p2 domain and express the transcription factor *lhx3* (Seredick et al., 2014). V2a cells are ipsilaterally projecting INs that drive swimming behavior in larval stages (Kimura et al., 2013), and could therefore contribute to early motor behavior. We used fluorescent in situ hybridization (FISH) to detect *lhx3* expression in wild-type and *fmr1* mutants, finding that expression was limited to cells in the ventral half of the spinal cord, including a small population of *lhx3*^+^ cells abutting the central canal that appeared to be KA neurons (white arrowheads, Fig. 3E,F), as well as ostensible V2a INs in the ventrolateral regions of the cord (yellow arrowheads, Fig. 3E,F). There was no change in the quantity of either *lhx3*^*+*^ subtype in *fmr1* mutants compared to wild-type (Fig. 3G,H). Taken together, our results indicate no clear changes in spinal glutamatergic neuron specification in *fmr1* mutants.

### Excess VeLD interneurons are produced in the absence of Fmrp

We used additional fate mapping approaches to determine the origin and identity of excess GABAergic INs in *fmr1* mutants. Based on the relative position of these cells in the cord, we reasoned they could derive from either the p2 or pMN domains (Park et al., 2004; Kimura et al., 2008; Andrzejczuk et al., 2018). We first tested the p2 origin hypothesis by examining gene expression specific to GABAergic V2b INs, which express the transcription factor *gata3* (Seredick et al., 2014). We detected *gata3* via FISH and then used IHC to label GABAergic neurons in both wild-type and *fmr1* mutants at 24 hpf, mainly finding *gata3*^*+*^GABA^+^ KA INs adjacent the central canal and a few *gata3*^*+*^GABA^+^ cells per section that were more proximal to the lateral edge of the cord (presumptive V2b INs, magenta arrowheads, Fig. 4A,B). There was no difference in *gata3*^+^GABA^+^ V2b cells (Fig. 4C) or total *gata3*^+^ cells (Fig. 4D) in *fmr1* embryos compared to wild-type controls. However, there were additional *gata3*^*-*^GABA^+^ cells in *fmr1* embryos (cyan arrowheads, B’), which suggests an alternative progenitor origin. This indicates that excess GABAergic INs in *fmr1* were not specified in the p2 domain.

**Figure 4.**
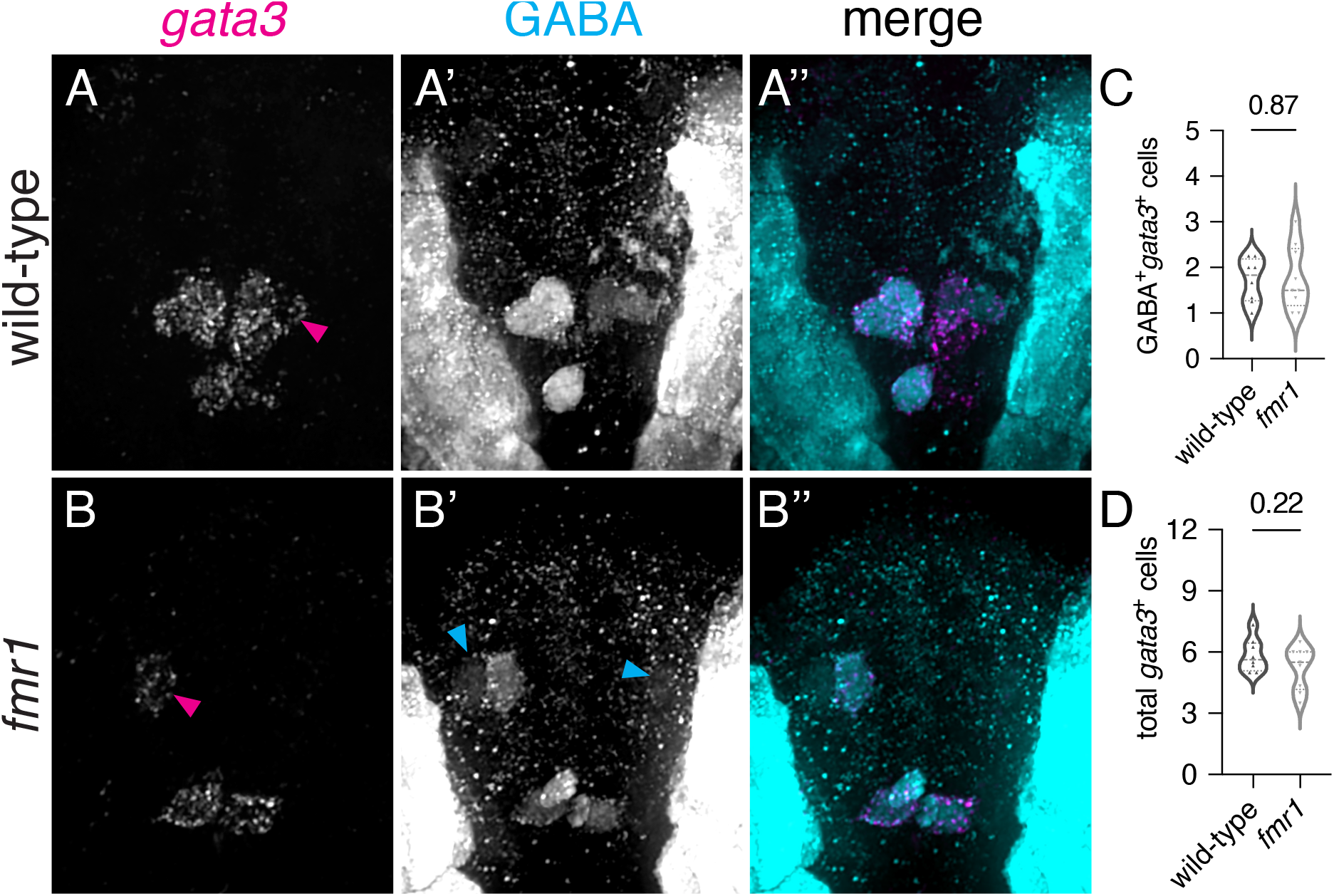
Excess GABAergic neurons in *fmr1* embryos are not specified in the p2 progenitor domain. Representative images showing immunohistochemistry for GABA in 24 hpf wild-type (A) and *fmr1* mutant embryos (B) alongside fluorescent RNA in situ hybridization to detect gata3 transcript (A’,B’). Quantification of GABA^+^*gata3*^+^ V2b cells (C) and total *gata3*^+^ cells at 24 hpf. GABA^+^*gata3*^+^ V2b neurons marked with magenta arrows (A,B), GABA^+^*gata*^+^ cells indicated with cyan arrows (B’). Scale bar = 10 μm.

As an alternative hypothesis for the origin of surplus GABA^+^ INs in *fmr1* mutants, we addressed IN specification from the pMN domain, which gives rise to a wide variety of cell types: cholinergic motor neurons, oligodendroglia, glutamatergic INs, a subpopulation of KA INs, and GABAergic ventrolateral descending (VeLD) INs (Park et al., 2004). We used IHC to detect GABA expression on trunk spinal cord sections of *olig2:*EGFP transgenic embryos, a pMN domain reporter (Fig. 5A,B). Although there was no change in pMN-derived GABA^+^ KA cells (Fig. 5C), there was a 71% increase in *olig2*^+^GABA^+^ cells (Fig. 5D) and a 35% increase in total GABAergic cells in *fmr1* embryos compared to wild-type controls. Moreover, presumptive VeLD INs in *fmr1* embryos were often located dorsally to bright EGFP^+^ pMN cells (arrowheads, Fig. 5B), whereas VeLD cells in wild-type were often masked by the reporter (Fig. 5A), which may suggest that Fmrp also regulates VeLD positioning in the cord. Taken together with our previous work, we can conclude that Fmrp regulates the specification of motor neurons, oligodendroglia, and VeLD INs from the pMN domain (Doll et al., 2021).

**Figure 5.**
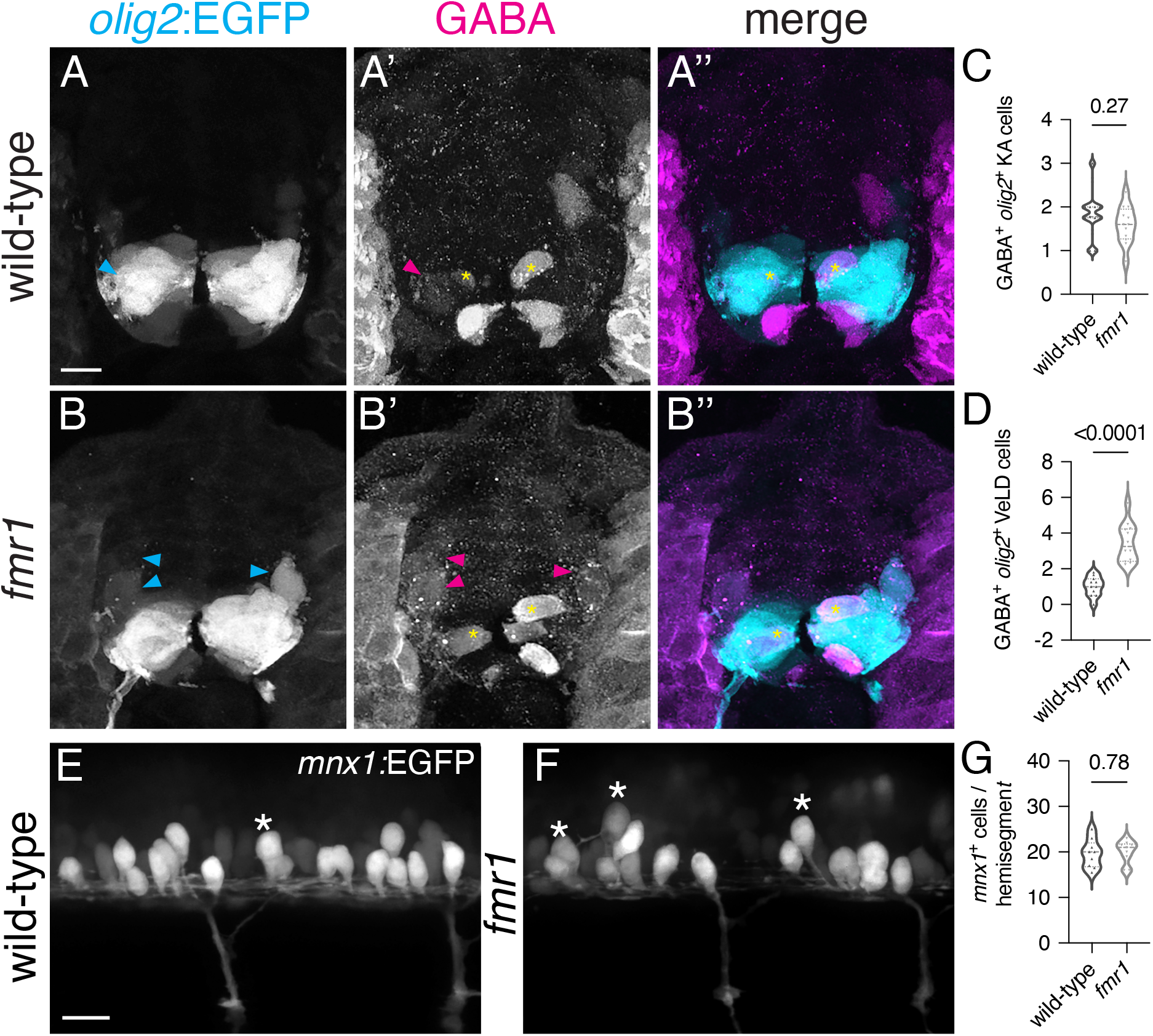
Surplus ventrolateral GABAergic interneurons in *fmr1* embryos are specified in the pMN progenitor domain. At 24 hours post-fertilization, there are two populations of pMN-derived GKBK^+^*olig2*^+^ cells in the ventral spinal cord, as shown through immunohistochemistry for GABA on transverse trunk spinal cord sections of Tg(*olig2*:EGFP) embryos: dorsal KA INs (yellow asterisks, A’,A”,B’,B”) and VeLD INs (cyan and magenta arrowheads, A,A’,B,B’). Quantification of KA INs (C), VeLD INs (D) in wild-type and *fmr1* embryos. VeLD INs in *fmr1* embryos are positioned more dorsally in the spinal cord (cyan arrowheads, B). (E,F) Live lateral images of trunk spinal cord from embryos expressing *mnxI*:EGFP, a reporter of primary motor neurons and VeLD interneurons. Asterisks indicate presumptive VeLD interneurons, with large soma situated dorsal and rostral to primary motor neurons. (G) Quantification of the average number of EGFP^+^ cells per hemisegement at 24 hpf. Scale bars = 10 μm.

Finally, we used the *mnx1:*EGFP transgenic reporter to provide both spatial and quantitative readouts of how Fmrp regulates the development and organization of early-born MNs and VeLD INs in living embryos (Flanagan-Steet et al., 2005). At 24 hpf, *mnx1:*EGFP is expressed in clusters of primary motor neurons and VeLD interneurons that are stereotypically arranged in each hemisegment of the spinal cord (Fig. 5E,F). VeLDs are positioned slightly dorsal to MNs at the rostral end of each hemisegment in the early embryonic cord of wild-type embryos, with large oval cell bodies (Fig. 5E; Seredick et al., 2012). In *fmr1* embryos, despite the increased presence of presumptive VeLD neurons (Fig. 5F), there was no change in the total quantity of *mnx1*^+^ cells in *fmr1* embryos compared to control (Fig. 5G). Taken together with our previous work showing reduced Isl1^+^ MNs in *fmr1* embryos (Doll et al., 2021), this suggests that Fmrp regulates the fate of pMN progeny without altering the quantity of these early-born neurons.

## Discussion

The embryonic zebrafish spinal cord contains a simple, well-mapped network of spinal neurons that act in concert to facilitate the progressive development of swimming motion (Kuwada et al., 1990; Eisen, 1991; Granato et al., 1996). Motor neuron function, as the output of the locomotive circuit, is clearly crucial to the process, yet primary motor neuron subunits are produced as discrete islands in individual hemisegments (Seredick et al., 2012). Integrative interneurons provide an orchestrating influence in the generation of complete movements by linking motor neurons throughout the spinal cord (Saint-Amant and Drapeau, 2001; Warp et al., 2012). We predict that the disproportionate production of VeLD INs in *fmr1* embryos may drive hyperexcitable motor behavior in this FXS model that persists in larval and adult stages (Kim et al., 2014; Shamay-Ramot et al., 2015). This builds upon a hypothesis that VeLD interneurons synchronize motor output in the zebrafish spinal cord to drive the earliest spontaneous movements (Warp et al., 2012). In the future we plan to directly test roles for VeLDs in the initiation of embryonic motor behavior.

We also show that excess GABA^+^ VeLD interneurons in *fmr1* zebrafish are associated with increased postsynaptic scaffold production at 24 hpf, indicative of accelerated synaptogenesis (Fig. 2). As Gephyrin expression is activity-dependent (Flores et al., 2015; Ravasenga et al., 2022), this appears to be in line with hyperexcitable motor output in FXS models. However, relative Gephyrin punctal quantity was comparable between genotypes a day later at 48 hpf, and at this stage the Gephyrin puncta in *fmr1* were significantly smaller than control. This may indicate that maturation of postsynaptic sites is hindered by the absence of Fmrp (Craig et al., 1996) or that terminal differentiation is inhibited in *fmr1* embryos. It is unclear whether surplus GABAergic neurons in *fmr1* directly influence premature scaffold production or whether it is a consequence of hyperexcitable FXS circuits at large (Kramvis et al., 2013; Ng et al., 2013; Kashima et al., 2017; Schaefer et al., 2017; Hu et al., 2020).

Interestingly, it does not appear that GABA plays a role in the earliest spontaneous activity that initiates just a few hours before 24 hpf, as this behavior is largely insensitive to GABAergic and glutamatergic antagonism (though glycine appears to exert some influence; Downes and Granato, 2006; Saint-Amant and Drapeau, 2001, 2000). It is unknown when GABA neurotransmission initiates in the cord and the relative weight of this signaling on embryonic motor function. These embryonic findings appear to contrast established circuits in FXS and autism, which often show deficits in GABAergic neurons and GABA signaling (D’Hulst et al., 2006; Kratovac and Corbin, 2013; Hashemi et al., 2017). However, GABA is unique in neurodevelopment, as it is capable of depolarizing target neurons due to chloride homeostasis in immature neurons (Rivera et al., 2005; Khazipov et al., 2015). Reduced GABAergic/glycinergic scaffold production in embryogenesis (Fig. 2) could also contribute to diminished inhibition after the polarity shift, thus contributing to persistent hyperexcitability seen in zebrafish and other FXS models (Kim et al., 2014; Shamay-Ramot et al., 2015). We also plan to assess chloride transporter expression in vivo, as the Nkcc1 chloride importer and Kcc2 chloride exporter mediate the GABA polarity shift and the genes encoding these transporters may be translationally regulated by FMRP (He et al., 2014).

Ultimately, FXS pathology is rooted in molecular mechanisms due to the loss of the RNA binding protein Fmrp. Our work shows another role for Fmrp in the specification of a unique and influential cell type from the pMN progenitor domain (Doll et al., 2021). Based on our data and previous work, it appears that early-born pMN progeny are disproportionately specified in *fmr1* embryos (Fig. 5), with reduced Isl1^+^ motor neurons and excess VeLD INs (Doll et al., 2021). In related studies, disruption of Isl2 function or loss of *islet1* also lead to changes in pMN cell fate, such that VeLD-like cells are overproduced at the expense of primary motor neurons (Segawa et al., 2001; Moreno and Ribera, 2014). We predict that Fmrp-associated changes in neural patterning drive these shifts in specification, leading to disproportionate ratios of early-born MNs and VeLD INs, as well as later-born oligodendrocyte lineage cells (Doll et al., 2021). In the future we will characterize the genetic signatures of pMN progenitors and identify Fmrp target mRNAs in pMN-derived cells to help reveal the mechanisms that underlie proportionate cell production during motor circuit development.

## Acknowledgements

The authors would like to thank Natalie Carey for insight on GABA; Angie Ribera, Melissa Wright, and Doug Hicks for transgenic lines; Bruce Appel and the Appel laboratory for support and feedback. This work was supported by the US National Institute of Health (NIH) grant R21 NS117886 to C.D.

